## Supplemental Tables and Figures for "The crystal structure of a simian Foamy Virus receptor binding domain provides clues about entry into host cells"

### Supplementary Tables and Figures

#### Table of Contents

***Supplementary Tables and Figures*** _____ ***1***

**Supplementary tables** _____ **2**

Table S1: X-ray crystallography data collection and refinement statistics. _____ 2
Table S2: Secondary structure content in GII RBD _____ 4
Table S3: Intramolecular interactions within GII RBD _____ 5

**Supplementary figures** _____ **7**

Figure S1: The fold of the FV RBD is maintained by hydrophobic and polar interactions _____ 7
Figure S2: Mobile loops decorate the apex of the RBD _____ 8
Figure S3: Comparison of glycosylated vs deglycosylated RBD structures _____ 9
Figure S4: Comparison of the SFV RBD fold with that of the RBD of Orthoretroviruses _____ 11
Figure S5: Sequence conservation of FV Env _____ 12
Figure S6: Intramolecular contacts between N8 sugar and RBD _____ 16
Figure S7: Functional features of FV EBD mapped onto the structure _____ 17
Figure S8: AlphaFold models of FV RBDs _____ 18
Figure S9: FV RBD common core excludes a large portion of the upper subdomain _____ 20
Figure S10: The inter-protomer RBD contacts formed by the upper domain loops show poor sequence conservation _____ 21
Figure S11: Recombinant RBD variants remain monomeric in solution _____ 22
Figure S12: Flow cytometry gating strategy _____ 23
Figure S13: Effect of mutations on FVV release and infectious titer _____ 25
Figure S14: Structural basis for RBDjoin region being dispensable for binding to cells _____ 26

**References** _____ **27**

#### Supplementary tables

##### Table S1: X-ray crystallography data collection and refinement statistics.

| | **SFV GII RBD$^D$ (native) data (PBD 8AEZ)** | **RBD$^D$ (derivative) data** | **SFV GII RBD$^G$ (native) data (PBD 8AIC)** |
|---|---|---|---|
| **Data collection** | | | |
| Wavelength | 0.9786 | 1.907 | 0.9786 |
| Space group | $P3_221$ | $P3_121$ | $P6_1$ |
| Cell dimensions | | | |
| *a*, *b*, *c* (Å) | 99.5, 99.5, 120.6 | 99.6, 99.6, 120.9 | 123.6, 123.6, 191.6 |
| α, β, γ (°) | 90, 90, 120 | 90, 90, 120 | 90, 90, 120 |
| Resolution range (Å) | 49.73 - 2.574 (2.666 - 2.574) [a] | 46.06 - 3.171 (3.284 - 3.171) | 46.73 - 2.8 (2.9 - 2.8) |
| Total reflections | 453419 (44048) | 238580 (24149) | 875971 (87278) |
| Unique reflections | 22363 (2191) | 12192 (1165) | 40619 (4041) |
| Completeness (%) | 99.91 (99.50) | 99.60 (96.75) | 99.58 (99.14) |
| Redundancy | 20.3 (20.1) | 19.6 (20.1) | 21.6 (21.6) |
| $R_{merge}$ [b] | 0.2051 (0.9362) | 0.1776 (0.9898) | 0.199 (1.705) |
| $R_{pim}$ | 0.04701 (0.2117) | 0.04098 (0.2256) | 0.04364 (0.3719) |
| $I/\sigma(I)$ | 10.10 (1.85) | 20.30 (5.19) | 13.35 (1.45) |
| $CC_{1/2}$ | 0.986 (0.846) | 0.998 (0.917) | 0.997 (0.689) |
| **Refinement** | | | |
| No. reflections | 22355 (2190) | / | 40615 (4040) |
| No. of reflections for $R_{free}$ [c] | 1052 (110) | / | 2111 (208) |
| $R_{work}/R_{free}$ [d] | 0.213/0.253 | / | 0.194/0.229 |
| No. non-hydrogen atoms | 2945 | / | 6086 |
| Macromolecules | 2711 | / | 5448 |
| Ligands | 203 | / | 531 |
| Water | 93 | / | 321 |
| **Mean B value (Å$^2$)** | | | |
| Protein and sugar | 73.88 | / | 65.44 |
| Ligand/Ion | 94.16 | / | 114.49 |
| Water | 60.68 | / | 55.18 |
| **R.m.s. deviations** | | | |
| Bond lengths (Å) | 0.012 | / | 0.004 |
| Bond angles (°) | 1.62 | / | 0.77 |

| | | | |
|---|---|---|---|
| Ramachandran favored/outliers (%) | 95.99/0.31 | / | 95.69/0.00 |

[a] Values in parentheses are for highest-resolution shell.
[b] $R_{merge} = \Sigma_{hkl} \Sigma_i |I_i(hkl) - \langle I(hkl) \rangle| / \Sigma_{hkl} \Sigma_i I_i(hkl)$, where $I_i(hkl)$ is the *i*th observation of reflection *hkl* and $\langle I(hkl) \rangle$ is the weighted average intensity for all observations *i* of reflection *hkl*.
[c] The free set represents a random 5% of reflections not included in refinement
[d] $R = \Sigma_{hkl}(||F_{obs}|-|F_{calc}||)/\Sigma_{hkl}|F_{obs}|$, where $|F_{obs}|$ and $|F_{calc}|$ are the observed and calculated structure factor amplitudes, respectively.

Table S2: Secondary structure content in GII RBD

| Domain | Helical† (%) | β-strands (%) | Other‡ (%) |
|---|---|---|---|
| RBD | 30 | 14 | 56 |
| Lower | 45 | 17 | 38 |
| Upper | 20 | 12 | 68 |

† - α- and $3_{10}$ helices
‡ - B, S, T, X

The secondary structure content was calculated using 2StrucCompare webserver [1] at https://2struccompare.cryst.bbk.ac.uk/index.php.

#### Table S3: Intramolecular interactions within GII RBD

The intramolecular interactions were analyzed by ProteinTools program https://proteintools.uni-bayreuth.de [2].

| **Van der Waals contacts in the SFV RBD** | | | |
|---|---|---|---|
| Cluster # | Area (Å$^2$) | # of residues | Location (domain) |
| 1 | 153 | 4 | Lower |
| 2 | 1002 | 22 | |
| 3 | 2543 | 51 | Lower + upper |
| 4 | 82 | 2 | Upper |
| 5 | 77 | 2 | |
| 6 | 86 | 2 | |

**Polar contacts in the SFV RBD**

Lower subdomain

| Cluster | Donor - Acceptor | Distance (Å) |
|---|---|---|
| 1 | HIS225-ND1 -- GLN222-OE1 | 3.3 |
| 2 | **ARG226-NH1 -- GLU337-OE1** | 3.1 |
| | **ARG226-NH2 -- GLU337-OE2** | 2.4 |
| 3 | HIS234-ND1 -- TYR323-OH | 3.1 |
| 4 | THR242-OG1 -- GLN492-OE1 | 3.4 |
| 5 | HIS314-ND1 -- THR313-OG1 | 2.9 |
| 6 | TYR327-OH -- ASP320-OD2 | 2.5 |
| 7 | ASN331-ND2 -- ASN336-OD1 | 3.4 |
| 8 | **LYS342-NZ -- GLU339-OE1** | 3.3 |
| | **ARG343-NE -- GLU339-OE2** | 2.8 |
| | **ARG343-NH2 -- GLU339-OE2** | 3.2 |
| 9 | ASN351-ND2 -- GLU502-OE1 | 2.5 |
| | TYR551-OH -- GLU502-OE2 | 2.5 |
| 10 | **LYS352-NZ -- GLU495-OE2** | 3.2 |
| | TYR497-OH -- GLU495-OE1 | 3.1 |
| 11 | ASN368-ND2 -- ASN373-OD1 | 3.0 |

Upper subdomain

| Cluster | Donor - Acceptor | Distance (Å) |
|---|---|---|
| 12 | GLN244-NE2 -- GLN491-OE1 | 2.6 |
| | GLN491-NE2 -- SER488-OG | 3.0 |
| 13 | TYR267-OH -- ASP468-OD2 | 2.9 |
| 14 | THR288-OG1 -- GLU442-OE1 | 2.9 |
| | THR288-OG1 -- GLU442-OE2 | 3.4 |

| | | |
|---|---|---|
| | TYR456-OH -- GLU442-OE2 | 3.2 |
| 15 | **ARG297-NH1 -- ASP402-OD2** | 2.9 |
| | **ARG297-NH2 -- ASP402-OD1** | 2.6 |
| | **ARG297-NH2 -- ASP402-OD2** | 2.9 |
| | SER397-OG -- ASP402-OD2 | 2.8 |
| 16 | **ARG372-NH1 -- ASP378-OD2** | 2.8 |
| | **ARG372-NH2 -- ASP378-OD1** | 2.9 |
| | **ARG382-NH1 -- ASP378-OD1** | 2.8 |
| | SER375-OG -- ASP378-OD2 | 2.9 |
| 17 | TRP399-NE1 -- ASP254-OD1 | 2.7 |
| 18 | THR406-OG1 -- ASN409-OD1 | 3.4 |
| 19 | **ARG407-NE -- GLU400-OE2** | 3.1 |
| 20 | **ARG433-NH2 – GLU464-OE2** | 2.7 |
| 21 | TRP435-NE1 -- ASN462-OD1 | 3.0 |
| 22 | **ARG436-NH2 – ASP254-OD1** | 2.6 |
| 23 | THR452-OG1 -- ASP450-OD1 | 2.9 |
| 24 | SER461-OG -- GLU439-OE2 | 2.5 |
| 25 | GLN482-NE2 -- SER473-OG | 3.1 |
| 26 | **ARG537-NE -- GLU384-OE1** | 3.0 |
| | ARG537-NH1 -- TYR269-OH | 3.4 |
| | **ARG537-NH2 -- GLU384-OE2** | 2.8 |
| | **ARG537-NH2 -- ASP274-OD2** | 2.8 |
| | TYR269-OH -- ASP274-OD1 | 2.6 |
| | TYR275-OH -- GLU384-OE2 | 2.6 |

#### Supplementary figures

##### Figure S1: The fold of the FV RBD is maintained by hydrophobic and polar interactions

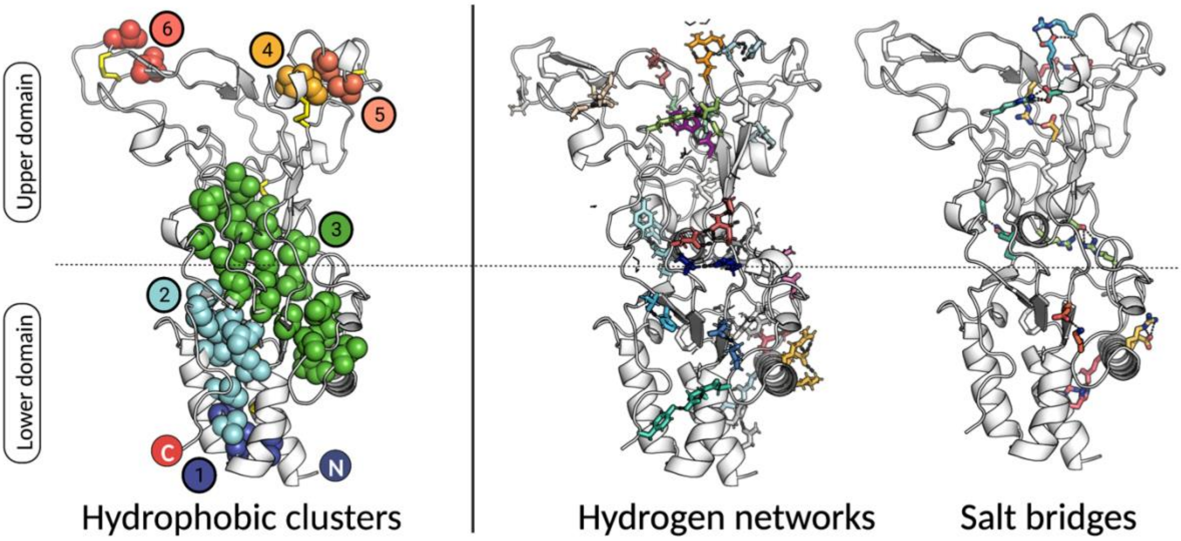


**Figure S1 legend:** The FV RBD core is formed by the hydrophobic residues grouped in 6 clusters - 2 in the lower subdomain (clusters #1 and #2), 3 in the upper subdomain (clusters #4, #5, #6), and the largest hydrophobic cluster (BSA=2451 Å$^2$ with 51 participating residues; cluster #3 shown in green) running in the direction of the longer axis of the RBD and containing residues from both domains. There are 24 networks of residues whose side chains contribute to 43 hydrogen bonds, with 21 charged residues forming 9 salt bridges. The area of the hydrophobic interfaces in the lower subdomain is about 6 times larger than in the upper subdomain, while the hydrogen bonds and salt bridges are more prevalent in the upper subdomain. The full list of intramolecular interactions and relevant details are given in Table S3. The intramolecular interactions were analyzed by ProteinTools program https://proteintools.uni-bayreuth.de [2].

#### Figure S2: Mobile loops decorate the apex of the RBD

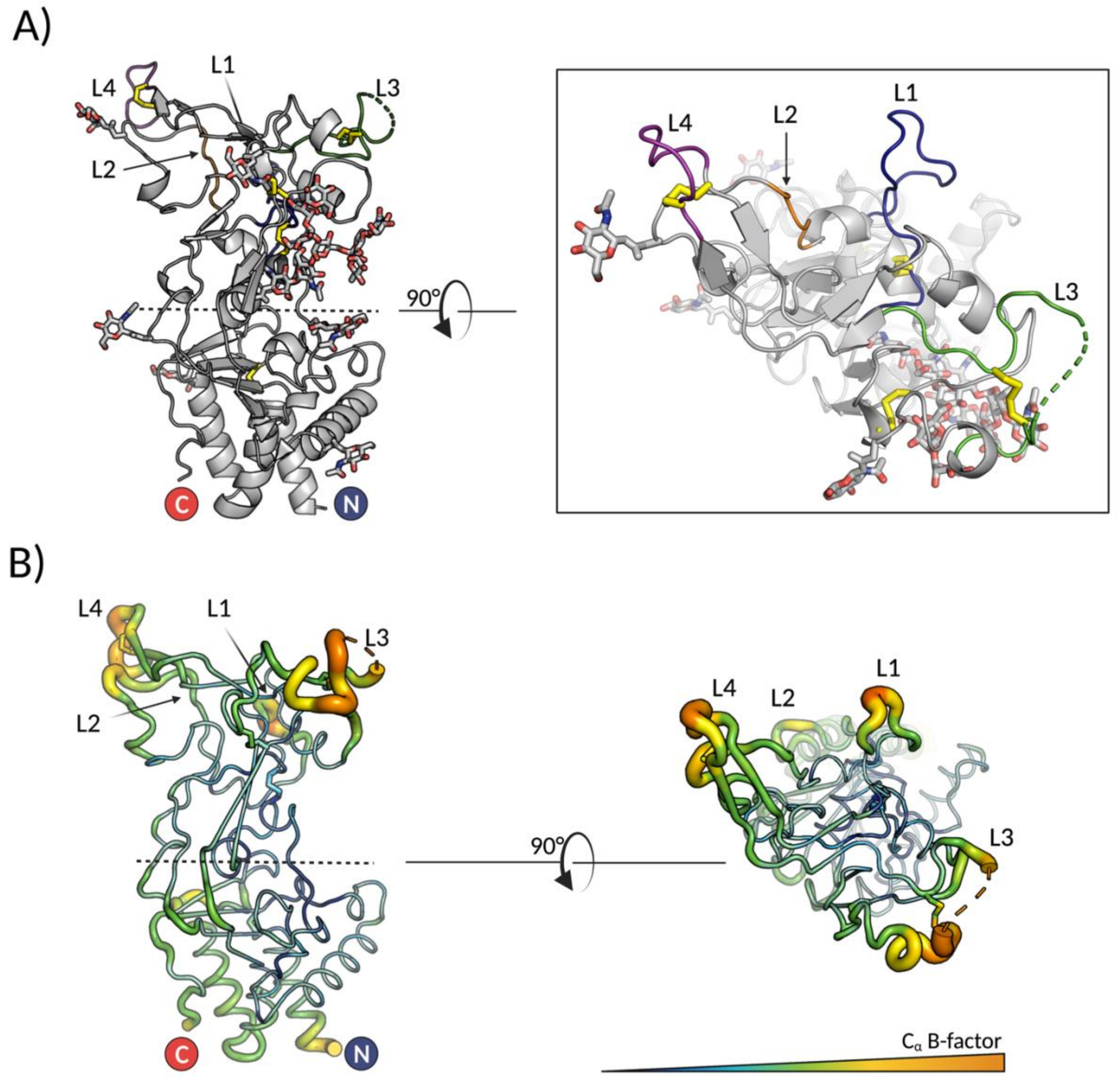


**Figure S2 legend: A)** The upper subdomains loops are designated as follows: loop 1 (L1, residues 253-270, connecting $\beta 2$ and $\eta 1$) in blue, loop 2 (L2, residues 276-281, connecting $\eta 1$ and $\beta 3$) in orange, loop 3 (L3, residues 414-436, connecting $\alpha 6$ and $\beta 9$) in green, and loop 4 (L4, residues 446-453, connecting $\beta 10$ and $\beta 11$) in dark purple. **B)** The RBD structure is shown in 'tube' presentation to illustrate the mobility, with the more flexible regions shown as thicker tubes. Coloring scheme corresponds to the $C_\alpha$ atomic B-factors (low to high B factors shown in blue to orange spectrum). The images were generated in Pymol [3].

#### Figure S3: Comparison of glycosylated vs deglycosylated RBD structures

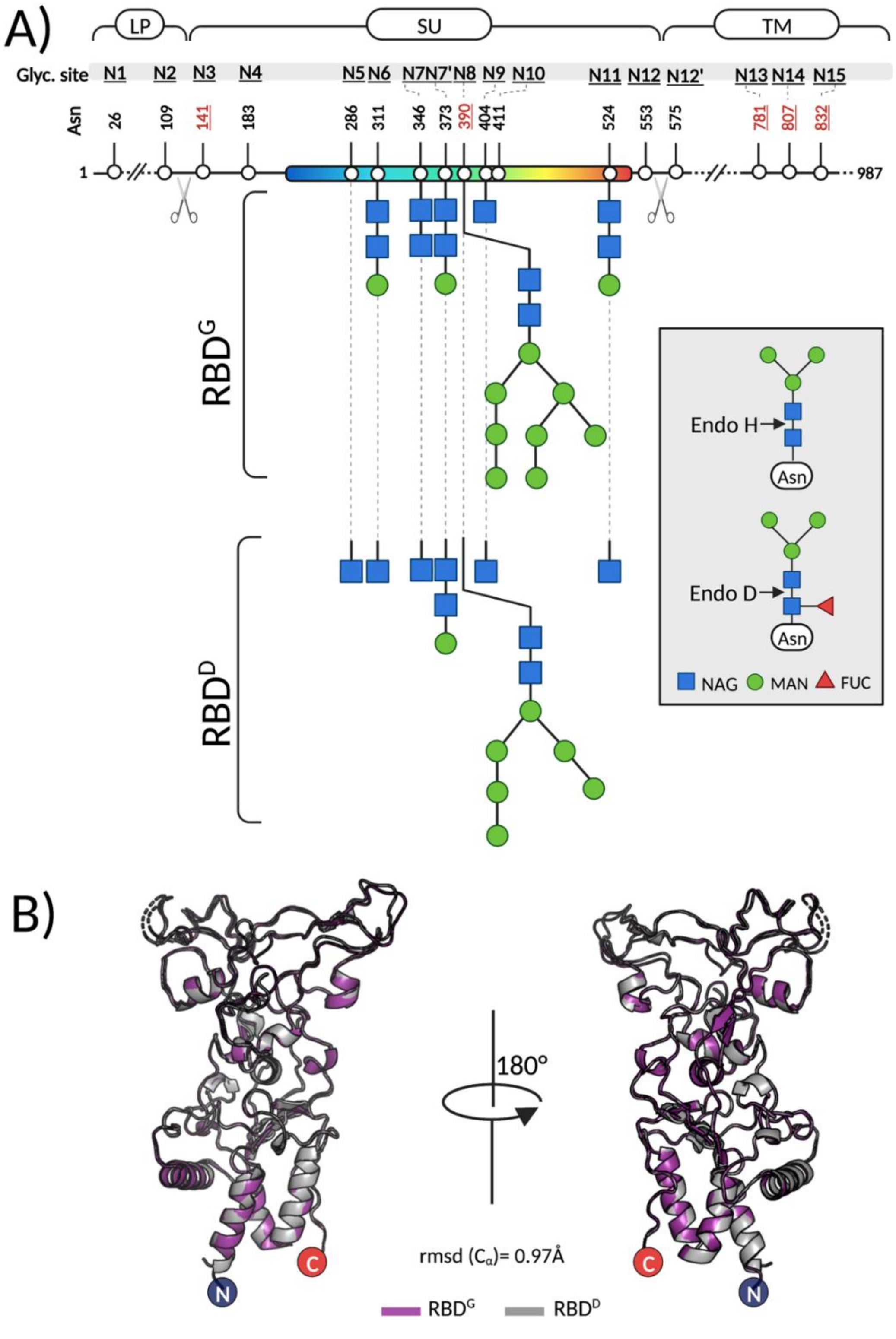


**Figure S3 legend: A)** Schematic representation of SFV Env and the 17 predicted N-glycosylation sites, labeled as N1 to N15 [4]. The sites that are 100% conserved in all FV Envs (Asn$^{141}$ (N3), Asn$^{390}$ (N8), Asn$^{781}$ (N13), Asn$^{807}$ (N14), Asn$^{832}$ (N15)) are indicated with red underscored letters. The two furin sites are represented by scissors. LP, SU and TM are the abbreviations for the leader peptide, surface subunit and transmembrane subunit, respectively. The sugar residues, N-acetyl glucosamine (NAG) and mannose (MAN) that could be resolved in RBD$^{D}$ or RBD$^{G}$ are shown.

The N-linked oligosaccharide core is shown in the grey inlet, with the cleavage sites indicated for the EndoD and EndoH glycosyades. A fraction of proteins expressed in insect cells contains an α1-6 fucose bound to the first NAG, rendering the sugar sensitive to cleavage by EndoD, but resistant to EndoH. Thus, both EndoD and EndoH were used for deglycosylation of the recombinant RBD. The figure was created in Biorender.com.

**B)** Superposition of the $RBD^G$ (purple) and $RBD^D$ (grey) structures done in Pymol [3].

Figure S4: Comparison of the SFV RBD fold with that of the RBD of Orthoretroviruses

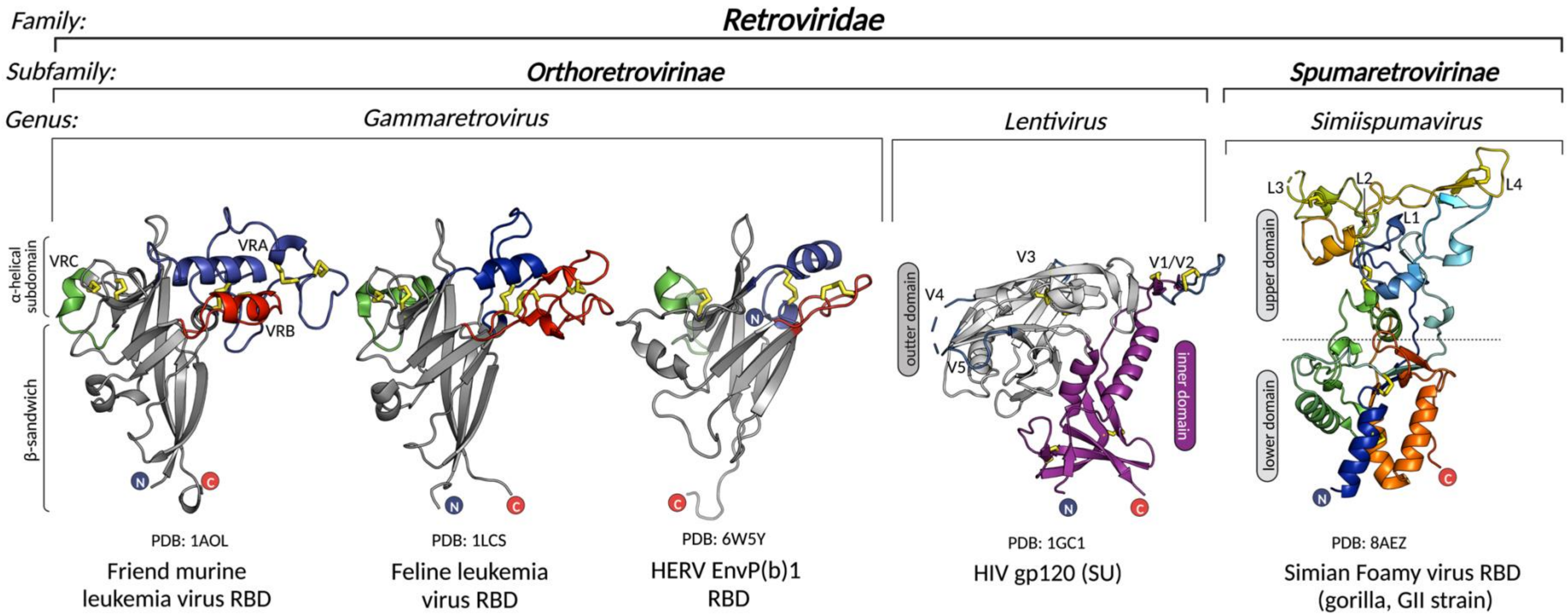


**Figure S4 legend:** Structures of RBDs from gammaretorviruses and of SU from HIV are shown to illustrate a lack of structural homology between the RBDs from different genera of retroviruses.

#### Figure S5: Sequence conservation of FV Env

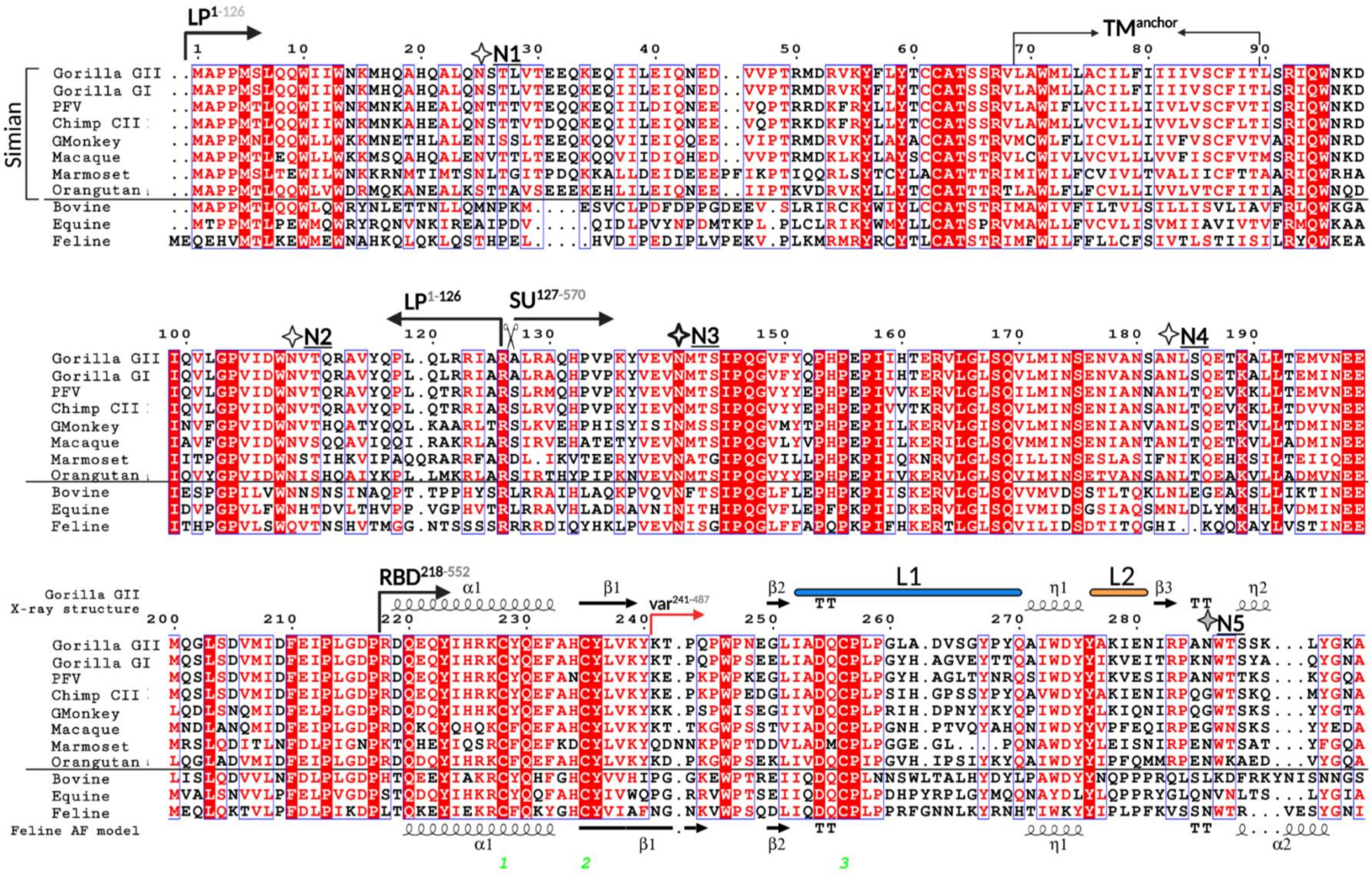

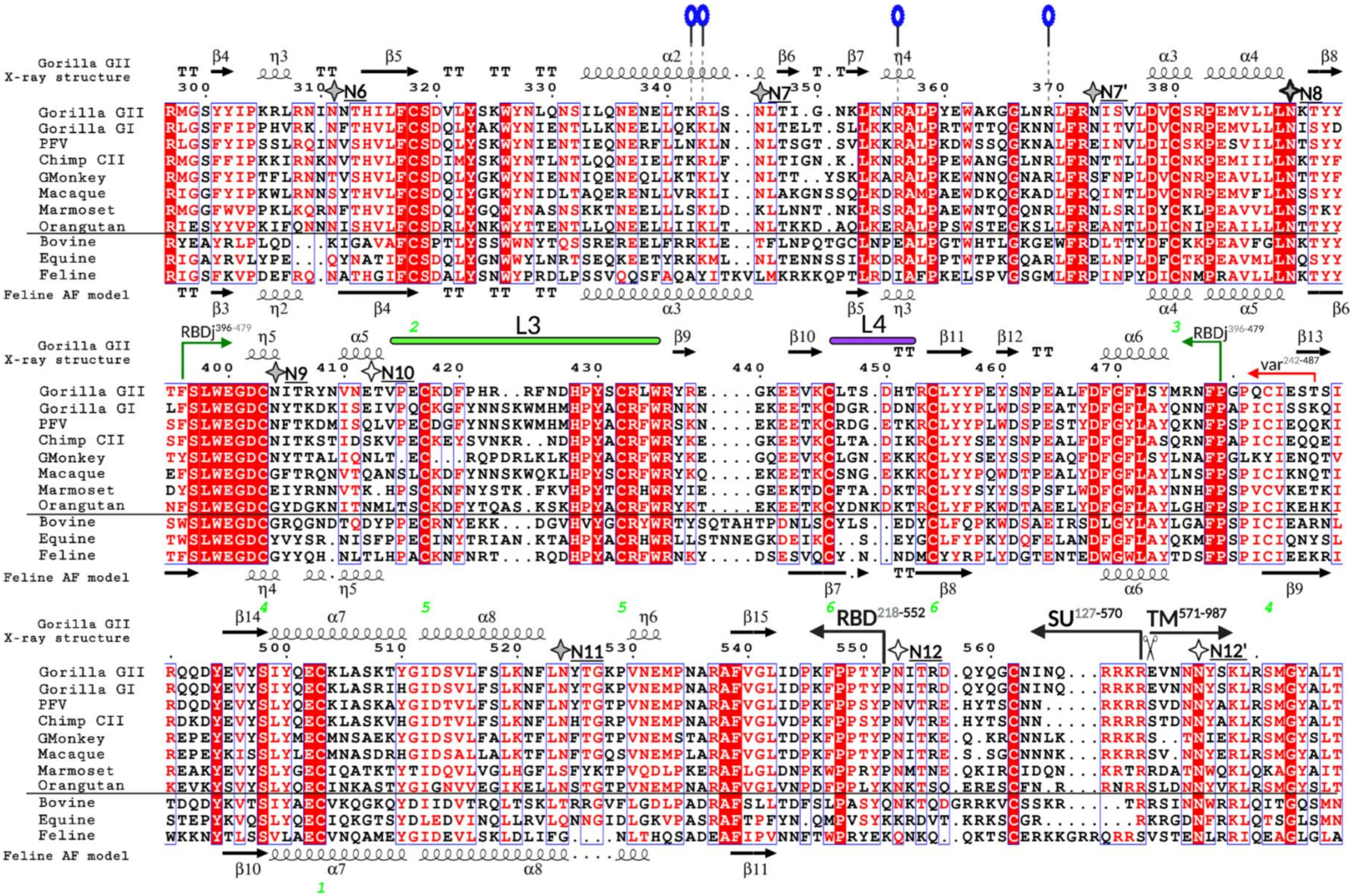
Gorilla GII X-ray structure
Gorilla GII
Gorilla GI
PFV
Chimp CII
GMonkey
Macaque
Marmoset
Orangutan
Bovine
Equine
Feline
Feline AF model
N6
N7
N7'
N8
N9
N10
N11
N12
N12'
L3
L4
RBDj396-479
var242-487
RBD218-552
SU127-570
TM571-987

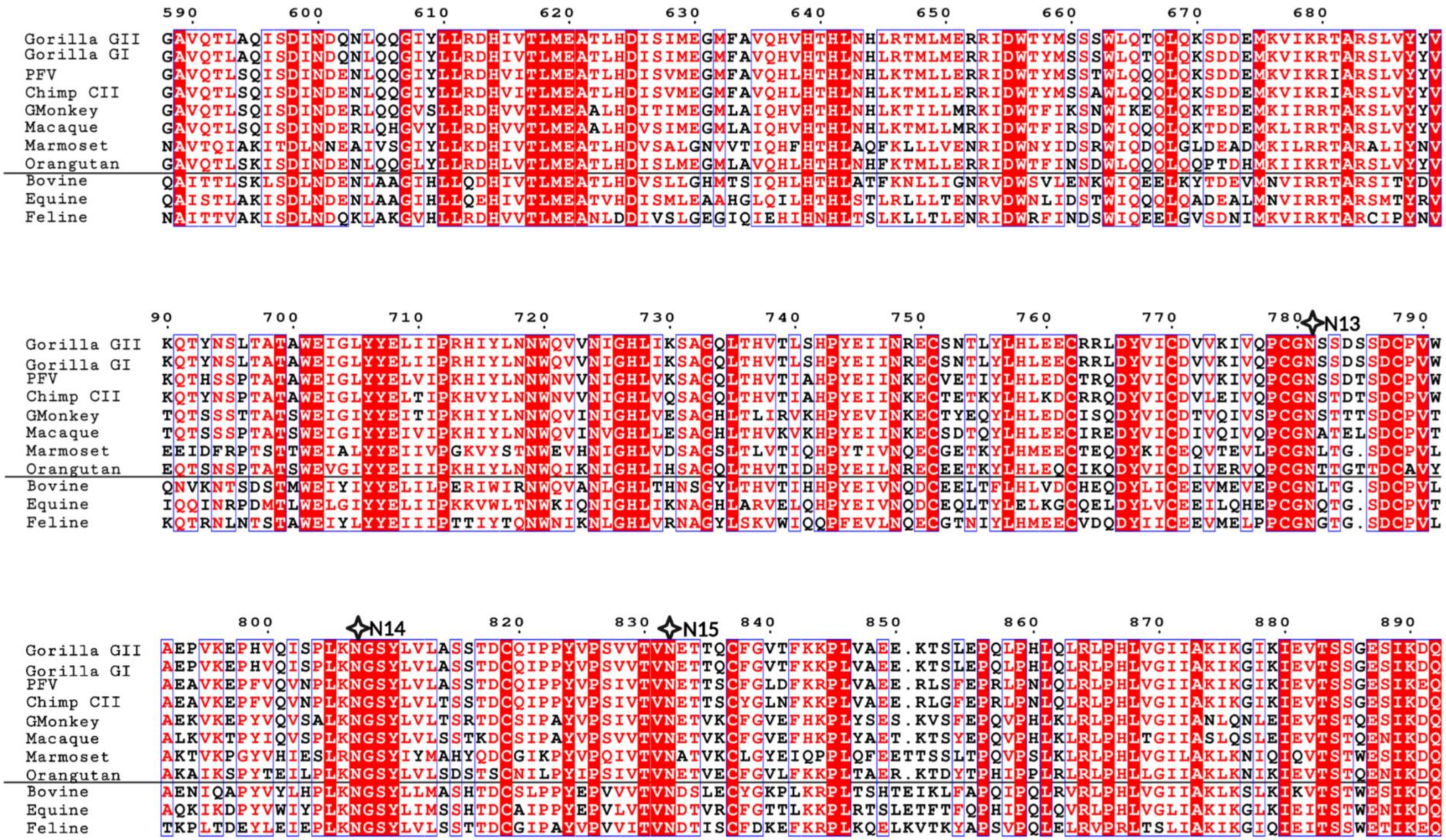
Gorilla GII
Gorilla GI
PFV
Chimp CII
GMonkey
Macaque
Marmoset
Orangutan
Bovine
Equine
Feline
N13
N14
N15

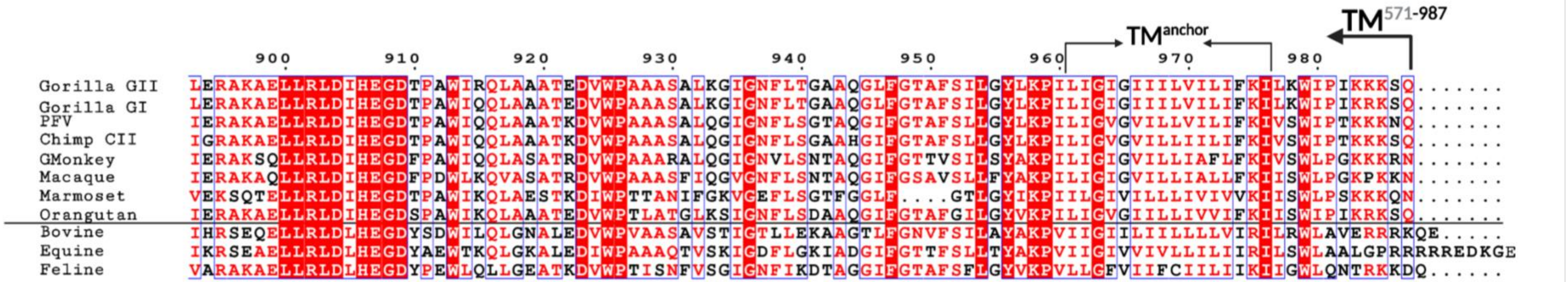


**Figure S5 legend**: Sequences corresponding to 11 FV Env were aligned in Clustal Omega [5] and the alignment was plotted using ESPript https://espript.ibcp.fr [6], with colors that indicate % identity (white letter, red background 100% identical; red letters, white background >70% identity; black letters, white background, <70% identity). The black, horizontal line separates simian from other FVs.

The secondary structure elements corresponding to the SFV RBD X-ray structure and the AF model for the feline FV RBD are plotted above and below the alignment, respectively. The N-linked glycosylation sites are indicated with stars. The already established nomenclature for the N-glycosylation sites (N1 to N15) is applied. The strictly conserved N-glycosylation sites have a thicker border, and the sites that carried sugars, which could be resolved in our structure, have grey filling. The residues interacting with heparan-sulfate are marked with blue ovals. Loops 1-4 are indicated with bars above the alignment and labeled as L1-L4, using the same color code as in Fig. S2. The boundaries for the LP, SU, RBD, TM subunit are shown, as well as for the RBD variable and RBDjoin regions (the numbering corresponds to that of gorilla SFV Env, GII-K74 genotype). To distinguish the TM subunit from the TM domain, which is the region spanning the membrane, the latter is referred to as the $TM^{anchor}$. The two furin sites are indicated with the scissors drawing.

The Env sequences used in the alignment were obtained from public databases and with following accession numbers: SFVggo_huBAK74 (GII-K74, genotype II gorilla SFV, GenBank: AFX98090.1), SFVggo_huBAD468 (GI-D468, genotype I gorilla SFV; GenBank: AFX98095.1), SFVpsc_huHSRV13 (CI-PFV, known as Prototype Foamy Virus genotype I chimpanzee SFV; GenBank: AQM52259.1), SFVpvePan2 (CII-SFV7, genotype II chimpanzee SFV; UniProtKB/Swiss-Prot: Q87041.1), SFVcae_LK3 (Genotype II African green monkey SFV; NCBI Reference: YP_001956723.2), SFVmcy_FV21 (genotype I macaque SFV; UniProtKB/Swiss-Prot: P23073.3), SFVcja_FXV (Marmoset FV; GenBank: GU356395.1), SFVppy_bella (Orangutan SFV; GenBank: CAD67563.1), BFVbta_BSV11 (Bovine FV; NCBI Reference: NP_044930.1), EFVeca_1 (Equine FV; GenBank: AAF64415.1), and FFVfca_FUV7 (Feline FV; UniProtKB/Swiss-Prot: O56861.1). Genotypes I and II have been defined for gorilla, chimpanzee, green monkey and macaque FVs [7] [8,9].

Figure S6: Intramolecular contacts between N8 sugar and RBD

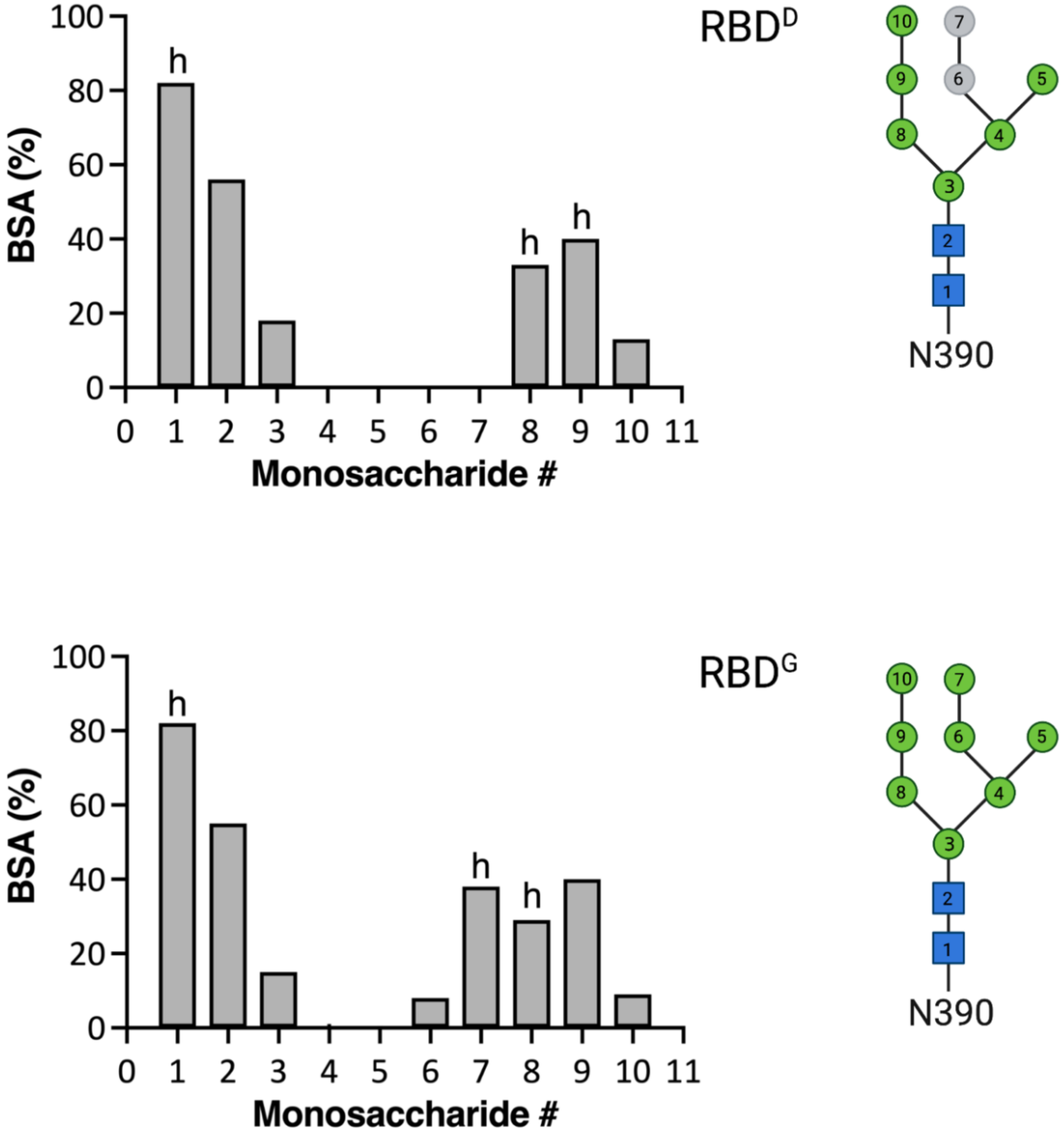


**Figure S6:** The buried surface area (BSA) for each sugar residue was calculated as a percent of the total surface area ($Å^2$) in ePISA [10] and plotted. Sugars that establish hydrogen bonds with the amino acids are indicated with letter ‘h’. Sugars 6 and 7 are colored in grey for RBD$^D$ because they were not resolved in the structure.

#### Figure S7: Functional features of FV EBD mapped onto the structure

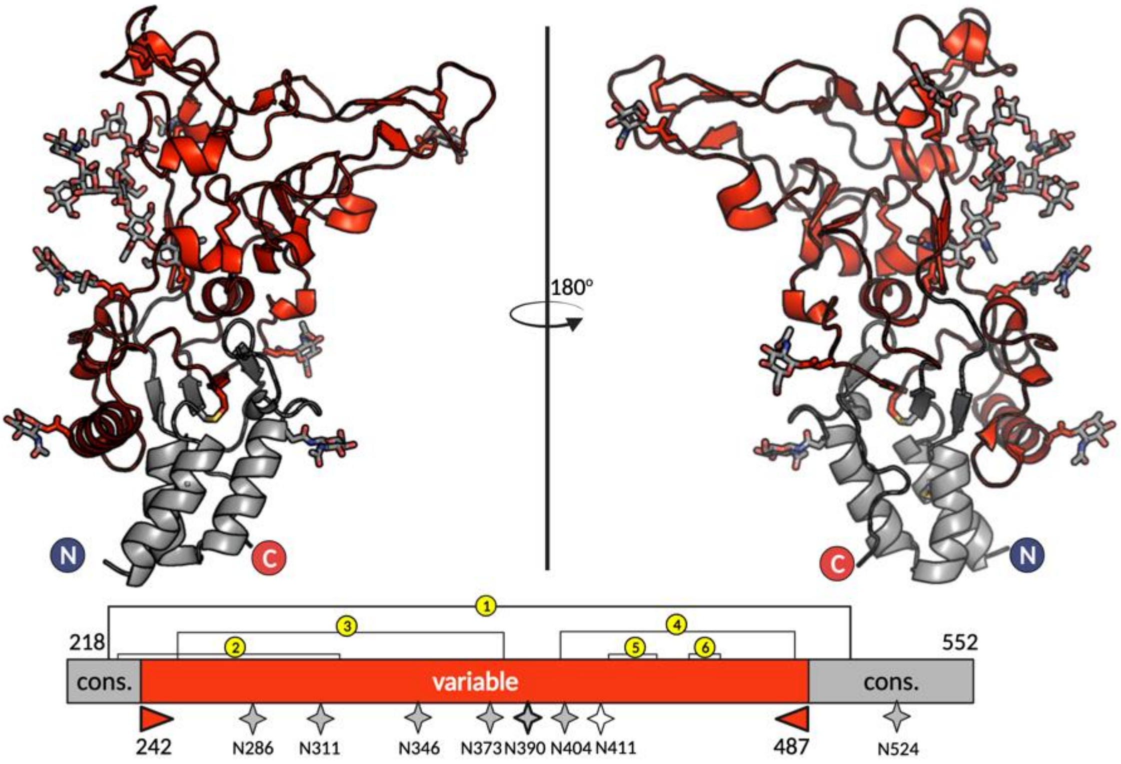


**Figure S7: Functional features plotted on the RBD structure.** The conserved ‘RBD$^{cons}$’ (residues 218-241 and 488-552) and variable regions ‘RBD$^{var}$’ (residues 242-487) are plotted on the X-ray structure of gorilla SFV RBD and colored in light grey and red, respectively. The glycosylation sites are indicated with the stars on the bottom.

#### Figure S8: AlphaFold models of FV RBDs

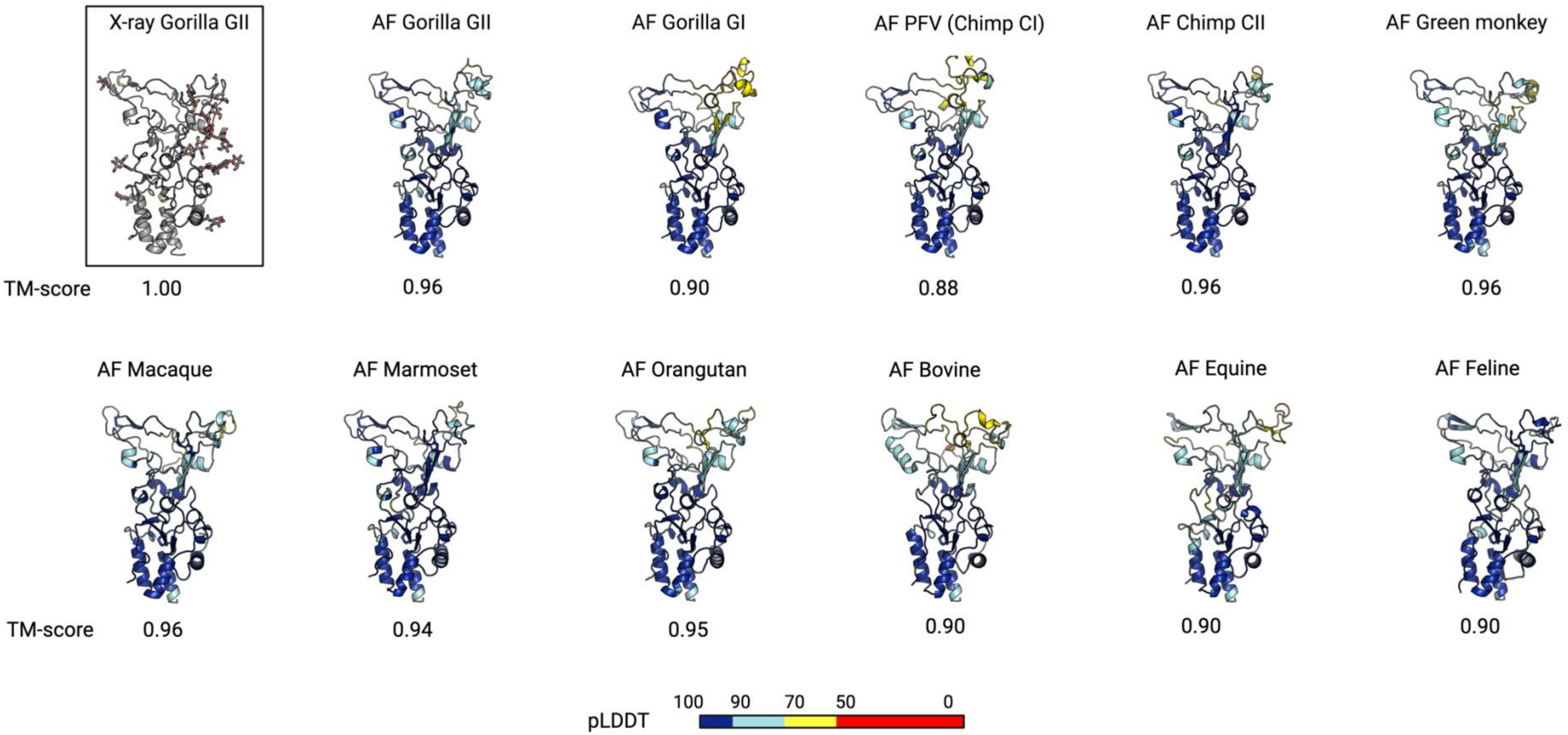


**Figure S8 legend:** Models generated by the AF prediction program [11] colored according to the per-residue confidence metric called 'predicted local distance difference test' (pLDDT). The pLDDT can have a value between 0 and 100, with the higher model confidence corresponding to the higher pLDDT number. pLDDT > 90 (rendered in blue on the panels) is the high accuracy cut-off, above which the backbone and rotamers are predicted with high confidence; values between 70 and 90 (cyan) correspond to the regions where the backbone conformation is correct; values between 50 and 70 (yellow) have low confidence and are not reliably predicted, and regions with pLDDT below 50 (red) should not be interpreted.

Structural superpositions of all AF models against each other were carried out using mTM-align server for multiple structural alignments[12,13] available at https://yanglab.nankai.edu.cn/mTM-align/. Below each model is a 'template modelling score' (TM-score), which is a length-independent scoring function reflecting the similarity of two structures[14]. The TM-scores can take values between 0 and 1, with the higher TM-score indicating higher structural similarity. The indicated TM-scores correspond to the pairwise superimposition of each AF model onto the X-ray Gorilla GII RBD structure.

#### Figure S9: FV RBD common core excludes a large portion of the upper subdomain

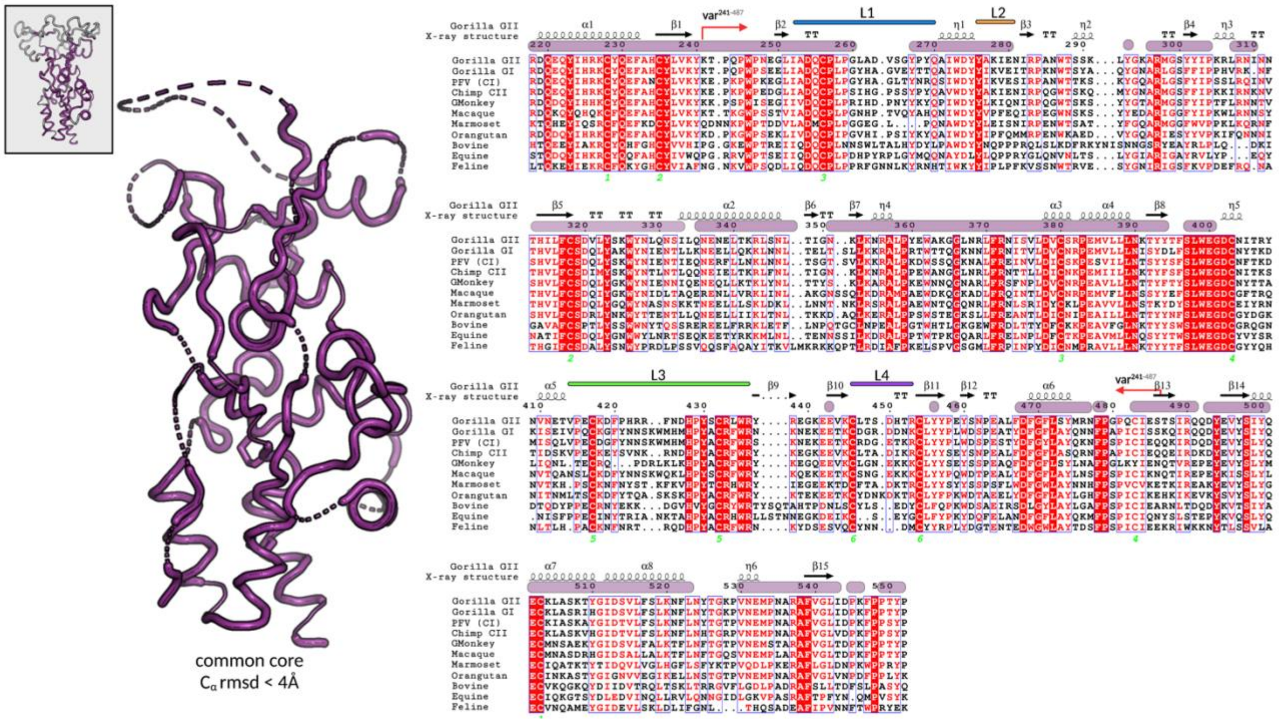


**Figure S9 legend**: Superposition of the RBD experimental structure and 11 AF models (Fig. S8) yielded a 'common core', model that includes the residues with C$\alpha$ rmsd < 4Å for all pairwise superpositions. Those residues are indicated with purple bars above the sequence alignment, which is colored using the same scheme as in Fig. S5. The small inlet in the upper left corner represents the entire RBD as a reference for comparison, with the common core colored in purple, and the remaining residues in grey. The structural and sequence alignments were carried out as explained in Figs. S8 and S5, respectively.

#### Figure S10: The inter-protomer RBD contacts formed by the upper domain loops show poor sequence conservation

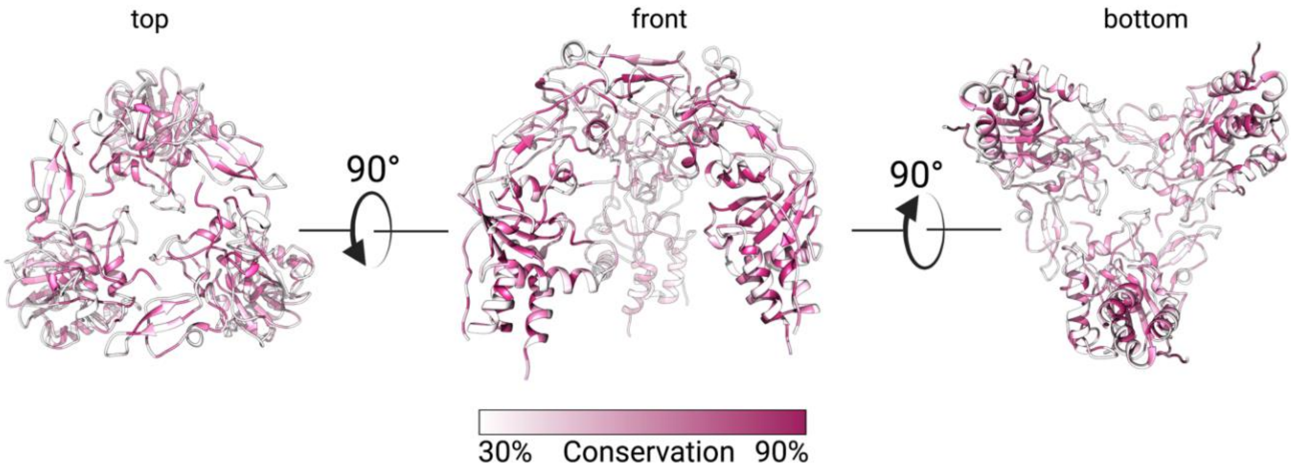


**Figure S10 legend:** Three SFV RBD protomers fitted in the electron density maps obtained for PFV Env [15], as shown in Fig. 4 are rendered by residue conservation. The % identity was calculated in Chimera [16] according to the sequence alignment shown in Fig. S5, and residues were colored with the white to maroon gradient as indicated with the color key. The residues showing less than 30% and more than 90% sequence identity are colored in solid white and maroon, respectively.

#### Figure S11: Recombinant RBD variants remain monomeric in solution

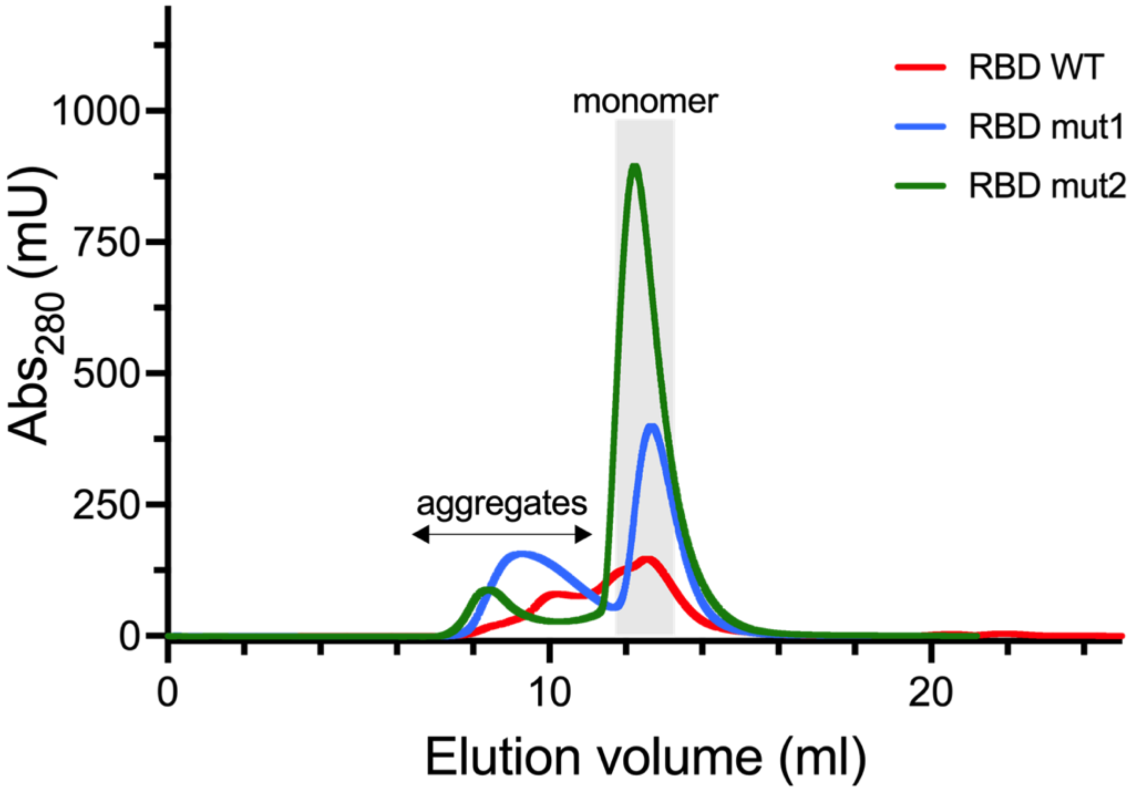


**Figure S11 legend**: The size exclusion chromatography (Superdex 200) profiles for the GII RBD expressed in mammalian cells are shown for the WT protein (red line) and the variants (blue and green lines).

#### Figure S12: Flow cytometry gating strategy

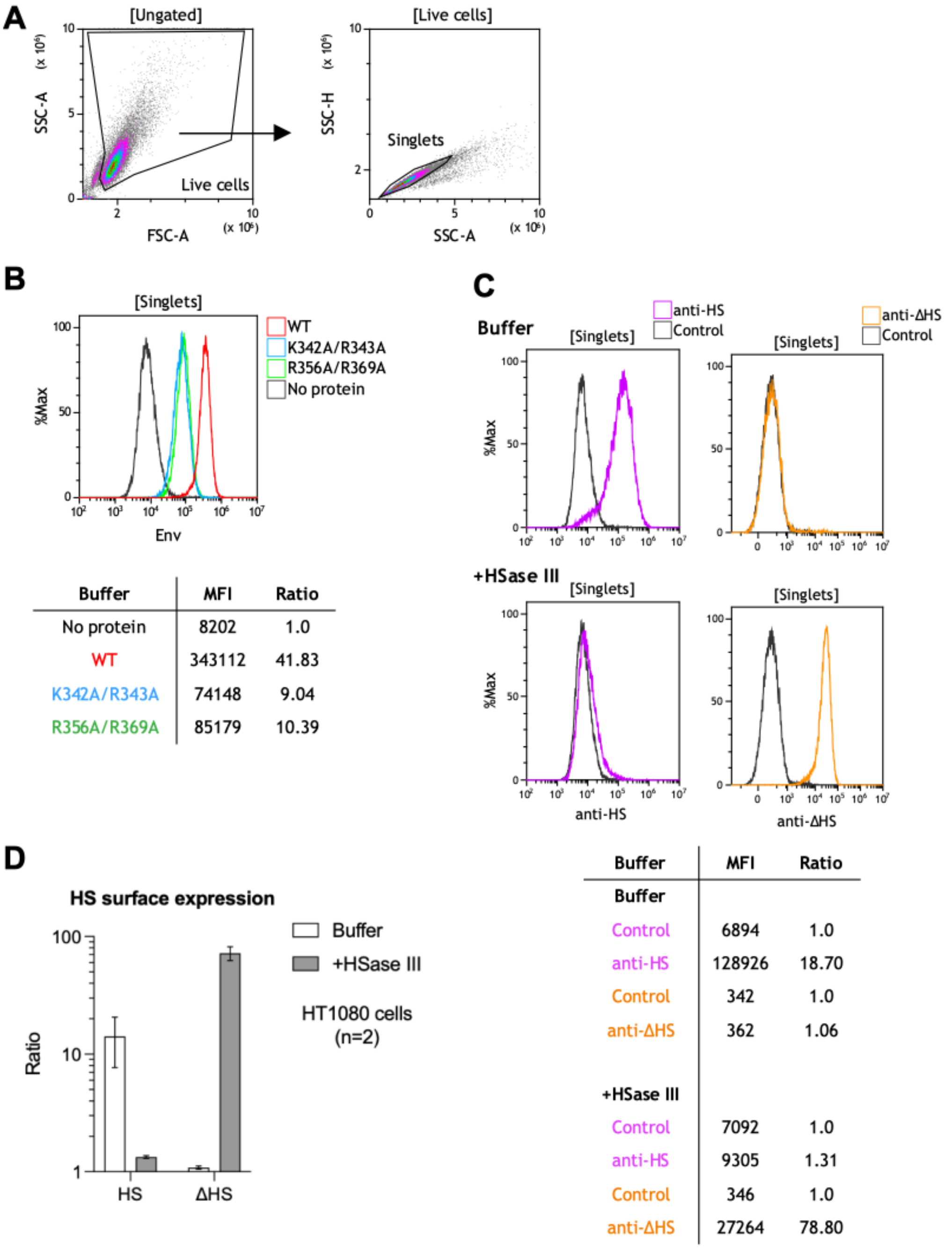


**Figure S12 legend**: Flow cytometry gating strategy for the detection of Env binding and HS expression on SFV-susceptible cell lines. Cells were treated with trypsin EDTA before labelling with Env proteins or anti-HS antibodies.

**A)** Representative example of HT1080 single cell selection: live cells were selected by a gate applied on an FSC-A/SSC-A dot-plot and a single cell gate applied on an SSC-A/SSC-H dot-plot.

**B)** Representative example of Env binding analysis. HT1080 cells were labelled with GII-K74 WT, K342A/R343A (mut1) or R356A/R369A (mut2) ectodomain proteins, anti-StrepMAB-Classic-HRP and anti-HRP-AF488 antibodies. Staining obtained on gated single cells are presented on the histogram overlay: MFI is presented on the x-axis and frequency is expressed as the normalized percentage of gated events on the y-axis (%Max). Cells labelled with secondary antibodies only (“control” condition, black curve) were used as a reference; Env-specific staining was quantified by the ratio of MFI from Env treated to untreated cells.

**C)** Representative example of heparan sulfate staining after treatment with heparinase III. HT1080 cells were treated with heparinase III or buffer and stained with the F58-10E4 antibody specific for heparan sulfate (anti-HS) and the F69-3G10 antibody specific for glycans exposed after heparan sulfate removal (anti-ΔHS). Staining obtained on gated single cells are presented on the histogram overlay: MFI is presented on the x-axis and frequency is expressed as the normalized percentage of gated events on the y-axis (%Max). Cells labelled with secondary antibodies only (“control” condition, black curve) were used as a reference; HS and ΔHS-specific staining was quantified by the ratio of MFI from labelled to control cells.

**D)** HT1080 cells were treated with heparinase III or buffer and stained with antibodies specific for HS (HS) or glycans exposed after heparan sulfate removal (ΔHS). Expression levels are calculated as the ratio of MFI from labelled to unlabeled cells (Fig. S12C). Mean and SD from two independent experiments are shown.

#### Figure S13: Effect of mutations on FVV release and infectious titer

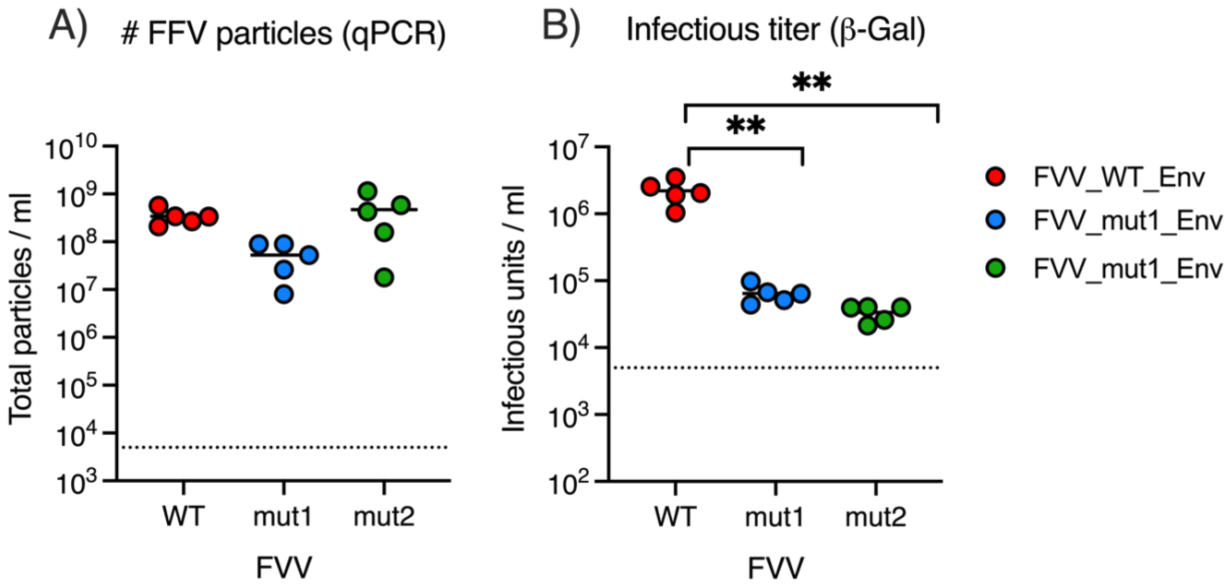


**Figure S13 legend**: Five batches of FVVs carrying wild-type GII-K74 SU (red), mut1 (blue) and mut2 (green) SU were produced, each represented with a single dot. **A)** The concentration of the vector particles was quantified by RT-qPCR of β-galactosidase transgene. Each batch was titrated twice, and mean titers are presented; lines represent mean values from the five FVV batches. The dotted line represents the quantification threshold.

**B)** FVVs infectious titers were quantified on BHK-21 cells. Each batch was titrated twice, and mean titers are presented; lines represent mean values from the five FVV batches. The dotted line represents the quantification threshold. The different FVVs were compared using the paired t test, * $p<0.05$, ** $p < 0.01$.

#### Figure S14: Structural basis for RBDjoin region being dispensable for binding to cells

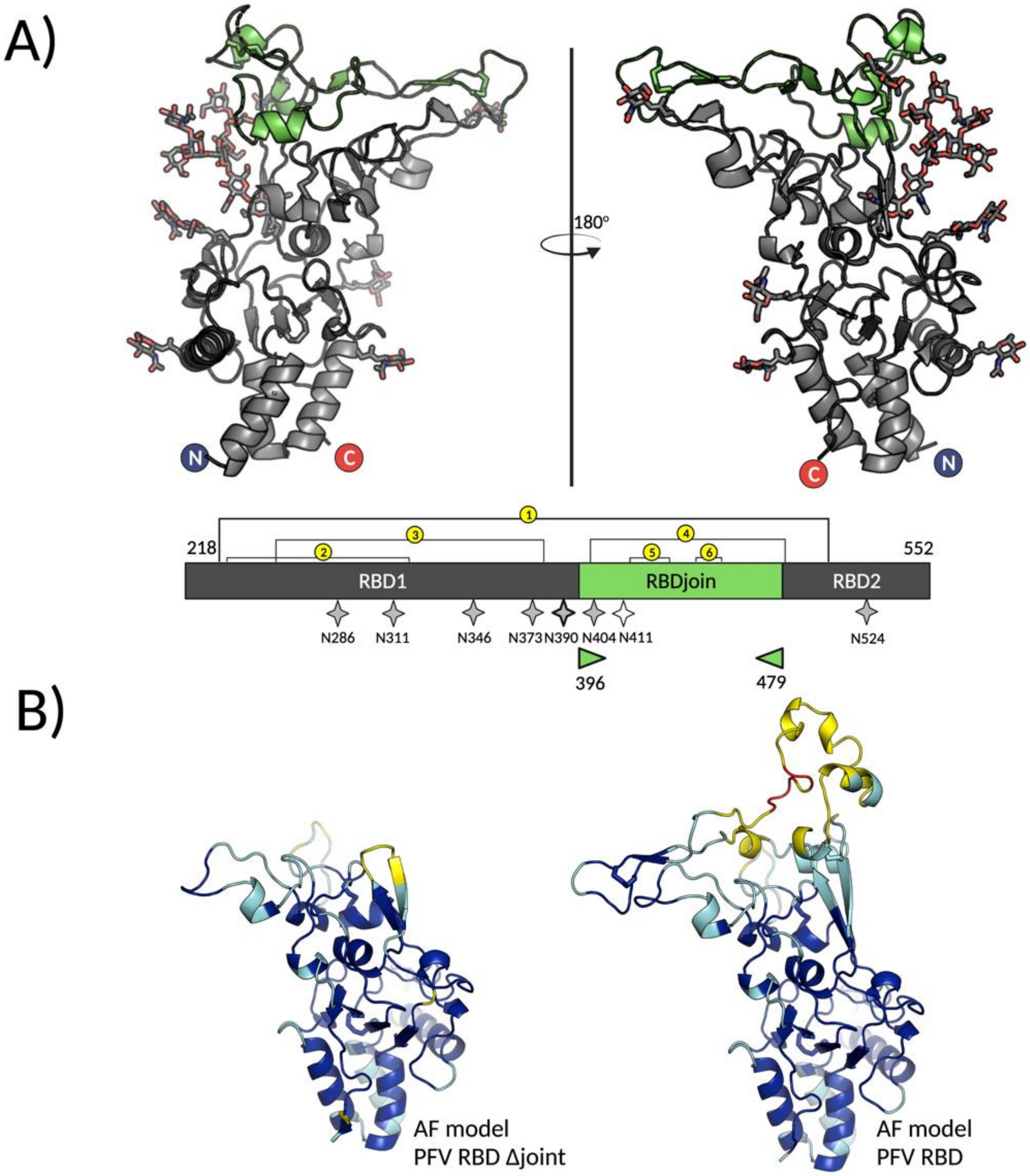


**Figure S14:** Functional features plotted on the RBD structure. **A)** The regions identified in the bipartite PFV RBD as essential [17] (indicated as RBD1 and RBD2 according to the more recent nomenclature [18]) and non-essential (or RBDjoin [18]) for SFV entry are colored in dark grey and green, respectively, and plotted on the X-ray structure of gorilla SFV RBD. The numbering corresponds to the gorilla GII RBD.
**B)** The AF models of the PFV RBD lacking the non-essential RBDjoin region (left panel) and of the whole PFV RBD (right panel) are colored according to the pLDDT values, using the same palette as in Fig. S8.
